# Linking extreme seasonality and gene expression in arctic marine protists

**DOI:** 10.1101/2021.11.11.467955

**Authors:** Magdalena Wutkowska, Anna Vader, Ramiro Logares, Eric Pelletier, Tove M. Gabrielsen

## Abstract

At high latitudes, strong seasonal differences in light availability affect marine organisms and restrict the timing of ecosystem processes. Marine protists are key players in Arctic aquatic ecosystems, yet little is known about their ecological roles over yearly cycles. This is especially true for the dark polar night period, which up until recently was assumed to be devoid of biological activity. A 12 million transcripts catalogue was built from 0.45-10 μm protist assemblages sampled over 13 months in a time series station in an arctic fjord in Svalbard. Community gene expression was correlated with seasonality, with light as the main driving factor. Transcript diversity and evenness were higher during polar night compared to polar day. Light-dependent functions had higher relative expression during polar day, except phototransduction. 64% of the most expressed genes could not be functionally annotated, yet up to 78% were identified in arctic samples from *Tara* Oceans, suggesting that arctic marine assemblages are distinct from those from other oceans. Our study increases understanding of the links between extreme seasonality and biological processes in pico- and nanoplanktonic protists. Our results set the ground for future monitoring studies investigating the seasonal impact of climate change on the communities of microbial eukaryotes in the High Arctic.

## Introduction

Solar radiation is a dominant energy source for life on Earth, and an important driver of evolution [1]. In the ocean, phytoplankton, mostly cyanobacteria and photosynthetic microbial eukaryotes, contribute half of the net primary production on Earth [2]. Light availability in the ocean declines with depth and forces a vertical distribution of species, with phototrophic organisms dwelling in the epipelagic zone (<200 m depth). The further from the equator, the more pronounced the annual changes in light regime, which at high latitudes is the strongest environmental driver of marine plankton phenology [3]. During the arctic polar night, the sun does not rise above the horizon for 4-6 months. The opposite happens during polar day, when the sun stays above the horizon for an equally long period. Extreme seasonality introduces profound limitations to biological processes in polar regions, and for centuries researchers perceived polar night as a period devoid of biological activity. Recent studies have reported substantial biological activity during the polar night; however, most of these studies focused on macroorganisms, predominantly zooplankton [4]–[6].

Our understanding of communities of marine microbial eukaryotes in the Arctic is primarily based on studies limited to a single sampling time point or cruises sampling along transects once or infrequently. However, disentangling the dynamics of changing community composition of organisms requires time series stations sampled at regular intervals [7], [8]. The world’s northernmost marine time series station (IsA) in Adventfjorden, Isfjorden, Svalbard (Figure 1), has been continuously sampled since December 2011 [9]. This endeavour generated metabarcoding-based knowledge regarding which marine microbial eukaryotes are present and active throughout the year [10], [11]. Seasonal dynamics of microbial eukaryotes can be analysed through many ecologically important measures and indices, such as diversity, biomass, cell counts, functions etc. In general, cell counts and biomass of microbial eukaryotes during polar night are lower compared to polar day, also at IsA [12], whereas OTU diversity is inversely proportional to this trend being higher during polar night [10]. However, proportion of plastid-bearing to heterotrophic cells is lower during polar night (reviewed in [4]).

**Figure 1.**
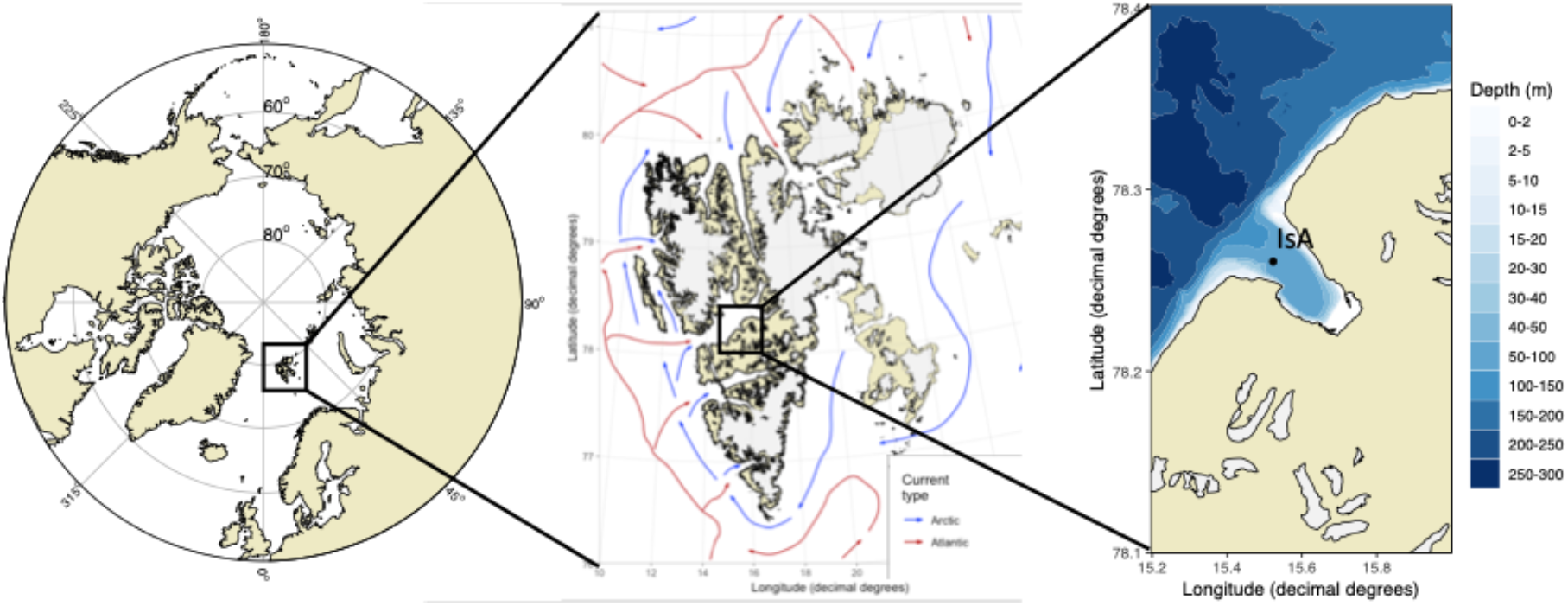
Location of the Isfjorden Adventfjorden (IsA) time series station in Svalbard.

Studies on the response of natural polar microbial communities to light/dark cycles are rare and cover a shorter timespan than the duration of the polar night [13]. Typical studies on dark survival of photosynthetic unicellular organisms are performed in laboratory conditions on single species cultures. Some of the key arctic microeukaryotic phototrophs were found ‘ribosomally active’ during polar night [10], [14]. Most of the primary production in the Arctic Ocean is performed by marine microbial eukaryotes when enough solar radiation is available [15], [16]. Outside this period these cells are assumed to use accumulated resources [17], decrease their metabolism [18], [19] or remain dormant [20]. However, many species of microbial eukaryotes instead of passively surviving prolonged darkness might switch their feeding strategy [21], [22], as is the case with mixotrophs [23].

Pico- and nanoeukaryotes play important roles in the marine environment, including photo-, heterotrophy or parasitism, with some species that can switch between these trophic modes [24]–[26]. These fractions of phytoplankton are challenging for classic microscopic taxonomy assignments or elucidating their roles. The analyses of their gene expression are especially helpful to understand what molecular processes they use to respond to environmental heterogeneity [24], [27]. Fluctuating environments might promote more stochasticity in gene expression of individual cells within populations, contribute to higher fitness and higher survival in the times of stressful conditions [28]. Nevertheless, community-level gene expression obtained by ‘omics’ methods was demonstrated as an effective predictor of current marine biogeochemical state [29]. In other words, the snapshot of metabolic functions performed by the community is tightly linked with environmental gradients present in the ecosystem at a given time.

We targeted the 0.45-10 μm size fraction of the microbial eukaryotic community from the IsA time series station to determine the dynamics of gene expression throughout a polar year, from which a 12 million eukaryote transcripts catalogue was built. Previous studies described higher diversity of microbial eukaryotes during polar night; thus, we hypothesize that the transcript diversity follows this trend. Given that light is the most important structuring force of community composition [3], we hypothesise that the light regime plays an essential role in controlling cellular processes in microbial eukaryotes. The presence of active phototrophic microbial eukaryotes during the polar night and their quick ecophysiological response to the return of light was confirmed by several studies [10], [13], [14], [30]. Hence, we hypothesise that genes involved in light-dependent processes, such as light-harvesting, are expressed also during polar night.

## MATERIALS AND METHODS

### Study site and sampling

The biological and environmental samples were collected at local noon at 11-time points between 14 December 2011 and 10 January 2013 from the Isfjorden Adventfjorden time series station (IsA); located on the west coast of Spitsbergen, Svalbard (N 78°15.6, E 15°31.8, Figure 1). At each of the 11 sampling dates, 30 l of seawater was collected from 25 m depth using a 10 l Niskin bottle (KC Denmark). Samples were kept in dark and cold conditions while prefiltered by gravity through 10 μm nylon mesh (KC Denmark) and then onto 8-12 47 mm 0.45 μm Durapore filters (Millipore) using a vacuum pump. Filters were fixed in 600 μl LB buffer (RNAqueous Total RNA Isolation Kit, Invitrogen, Thermo Fisher Scientific) 5-20 min after sampling, and subsequently flash-frozen in liquid nitrogen and stored at −80°C. At each sampling date, an 85m-depth vertical profile of environmental variables was obtained using a handheld SAIV 204 STD/CTD probe. Photosynthetically active radiation (PAR), size-fractionated chlorophyll *a* and nutrient concentrations (nitrate/nitrite, phosphate, silicate), were obtained as described in [10].

### mRNA extraction and amplification

Total RNA was extracted with the RNAqueous Total RNA Isolation Kit (Invitrogen, Thermo Fisher Scientific) according to manufacturer’s recommendation. Samples were thawed on ice, vortexed and kept on ice during RNA extraction. The thawed lysate was added to a tube with 200 μm molecular biology grade zirconium beads from pre-filled tubes. Extracts from filters collected at the same day were pooled together. We removed DNA using TURBO DNA-free Kit (Invitrogen, Thermo Fisher Scientific).

To test for the presence of PCR inhibitors, we used reverse transcription reaction of DNase-treated RNA samples. First, we denatured RNA molecules by incubating at 65°C for 5 min. in a mix of 1 μl of DNase-treated RNA, 1 μl of Random Hexamer Primer (at 100 μM concentration, Invitrogen, Thermo Fisher Scientific), 1 μl of dNTP mix (10 mM concentration each). Then we synthesised cDNA within reactions containing 4 μl 5x First Strand Buffer, 1 μl 0.1 M DTT, 1 μl RNase inhibitor (RNaseOUT, 40U/μl, Invitrogen, Thermo Fisher Scientific), 1 μl of SuperScript III Reverse Transcriptase (200U/μl, Invitrogen, Thermo Fisher Scientific) and 13 μl of denatured RNA samples. This reaction was incubated first for 5 min at 25°C, then for 45 min at 50°C and finally for 15 min at 70°C to inactivate reverse transcriptase. Amplification inhibitors in DNase-treated samples were removed by precipitation in 5 M ammonium acetate and absolute ethanol, using glycogen as an RNA carrier. RNA was amplified using the MessageAMP II aRNA Amplification Kit (Invitrogen, Thermo Fisher Scientific) according to manufacturer’s recommendations, extending the in vitro transcription step to 14 h. Amplified samples were dissolved in 100 μl of nuclease-free water and frozen at −80°C. Amplified mRNA was sent to GATC (Constance, Germany) where the libraries were prepared and sequenced on Illumina HiSeq 2500/4000, producing 150 bp paired-end reads.

### Data processing

Generated sequences were processed in four main steps: pre-processing, metatranscriptome co-assembly, mapping of reads from individual metatranscriptomes onto the assembly and finally annotation of assembled transcript isoforms (Figure 2). The quality of the data was assessed with FastQC v.0.11.5 [31]. The pre-processing step aimed to remove unwanted sequences from the metatranscriptomes. First, Illumina adapters were removed using BBDuk v. 37.36 [32]. Overrepresented sequences in each metatranscriptome reported by FastQC, consisting predominantly of poly(A) and poly(T) fragments, were removed with BBDuk. The same software was used to remove PhiX control reads. Although we used poly(A) selection to capture only eukaryotic mRNA during sample preparation, rRNA may remain in the samples [33]. Thus, we used SortMeRNA 2.0 [34] to remove sequences that mapped to rRNA. Lastly, BBDuk was used to remove sequences of quality score <20 and read length <25 bp (because the next step by default uses k-mers of that length).

**Figure 2.**
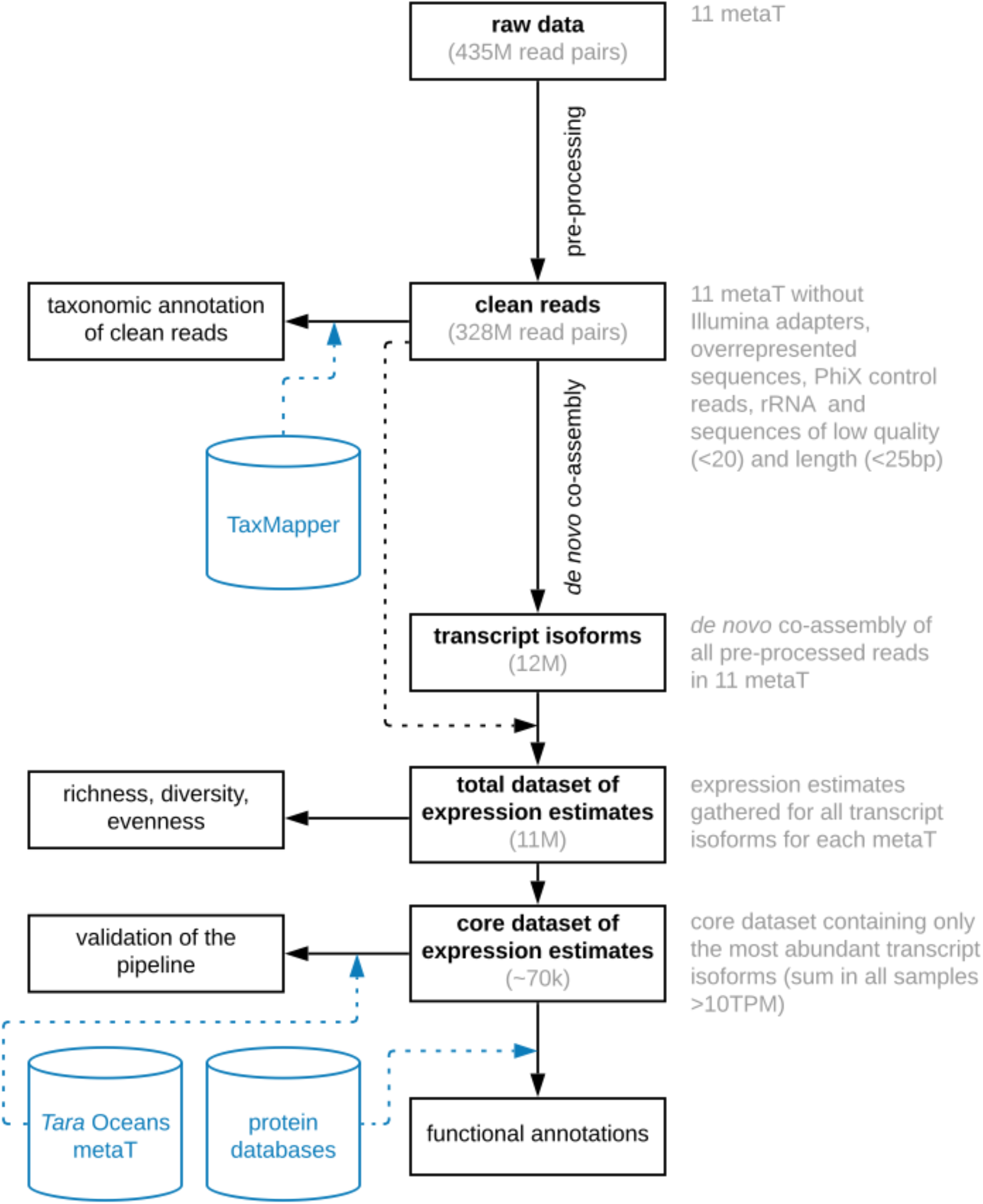
Sequencing data processing workflow. A separate metatranscriptome (metaT) was generated from each sample.

Detailed statistics on the initial library size of each metatranscriptome, and its change after each of the pre-processing steps can be found in Supplement 1. The 11 metatranscriptomes, containing jointly ~328 million read pairs, were *de novo* co-assembled into a gene catalogue using Trinity [35], [36]. Digital normalization removed 10 million read pairs with a median k-mer abundance of <2 (--min_cov 2) and >50 (--max_cov 50) prior to the co-assembly. The initial assembly step of Trinity – *Inchworm*, ran on 199 million of all read pairs with no further normalisation. The assembled output ran through the remaining part of the co-assembly, first constructing de Bruijn graphs (*Chrysalis* step) and then resolving them (*Butterfly* step). Expression levels were estimated by mapping clean reads against the gene catalogue in RSEM 1.3.0 [37]. Due to varying numbers of reads in each of the metatranscriptomes (Supplement 2) and to assure between-sample comparison [38] we used a relative measure of transcripts per million reads (TPM).

### Annotations

*De novo* assembly produced 12 245 433 transcript isoforms, with clean reads mapping at least once to 11 010 859 isoforms. Most transcripts were characterized by low sum of relative abundance across samples (8 transcripts with > 10 000 TPM, 154 with > 1000 TPM, 3483 with > 100 TPM, 68 166 with > 10 TPM and 2 390 862 with > 1 TPM; Supplementary 3). To increase the robustness of analyses and avoid stochasticity due to low abundance transcripts, further analyses were carried out on a core dataset that contained 68 166 of the most abundant transcript isoforms for which the sum of TPM in all the samples was greater than 10 (Figure 2; from now on we will refer to the transcript isoforms as “transcripts”). Coding regions were predicted using TransDecoder 5.1.0 (https://github.com/TransDecoder/TransDecoder/). The core dataset was functionally annotated using Trinotate 3.3.1 with default parameters [39]. Similarities between the *de novo* assembled transcripts/predicted coding regions and proteins in the UniProt database [40] were assessed using blastx/blastp, with max_target_seq = 1 and e-value 1e-3 (BLAST+) [41]. Protein domains were identified with HMMER3 [42] against the Pfam database (31.0 release) [43]. Functional annotations were retrieved with Trinotate based on blast results against Pfam and protein domains identified using eggNOG 3.0 [44], The Gene Ontology (GO) [45] and Kyoto Encyclopedia of Genes and Genomes (KEGG) [46], [47]. We focused on the most abundant GO terms dataset corresponding to biological processes, molecular functions, and cellular compartments with an arbitrary value of > 5000 TPM for each GO term.

Taxonomy was assigned to clean reads using the TaxMapper search tool and corresponding database with default settings [48]. Reads were mapped to two taxonomic levels: seven main eukaryotic lineages (supergroups, e.g., Alveolata) and 28 groups within these lineages (e.g., Apicomplexa, Chromerida, Ciliophora, Dinophyta and Perkinsea within the Alveolata supergroup).

To validate the process of the *de novo* assembly, we mapped transcripts in our core dataset against metatranscriptomic data from the *Tara* Oceans expeditions, including *Tara* Oceans Polar Circle sampled in 2013. The reads mapping pipeline used is the same as described previously [49]. Briefly, reads from each *Tara* Oceans’ metatranscriptomic read set were mapped onto transcript isoforms in our core dataset using bwa [50] and 95% identity over at least 80% of the length of the read picking the best match (or in case of several putative best matches – a random one).

### Statistical methods

All statistical analyses were performed in R v3.5.2 [51], and data were visualised using *tidyverse* v1.2.1 [52]. Principal component analysis (PCA) was calculated on centred and scaled data with *prcomp* function (*stats* package v3.5.3) and visualised using *factoextra* v1.0.5. To explore differences between transcript abundances a Bray-Curtis dissimilarity matrix (*vegdist* function in *vegan* package v2.5-4 [53]) was constructed and clustered using a ‘complete’ method within *hclust* function (*stats* package v3.5.3). *Pvclust* package was used to assign support to the clustering topology [54]. To identify the strongest contribution of individual transcript isoforms to clustering patterns, we applied the *simper* function on the transcript matrix.

To explore GO annotations, for each metatranscriptome, we summarised relative counts for each transcript that was assigned to a specific GO term. We explored each of the three categories of GO terms: molecular functions, biological processes, and cellular components. For each category, a Bray-Curtis dissimilarity matrix of GO abundance tables was used to calculate global non-metric multidimensional scaling (GNMDS [55]). The *envfit* function (*vegan* package) was used to fit environmental parameters onto the GNMDS ordination. Analysis of similarities (*ANOSIM*; *vegan* package) was used to test if there were differences between polar day and polar night associated with light. The simper function (*vegan* package) was used with 999 permutations to elucidate GO terms that contributed the most to the difference between polar night and polar day within the three GO categories. In this analysis, the September sample was excluded due to being from a time of mixed light conditions in the transition between polar day and polar night. *Simper* analysis identified GO terms that differed between polar day and polar night. These terms were then called “overrepresented” if the differences in means were statistically significant. Subsequently, all GO annotations within biological processes containing the words ‘light’ or ‘photo’ (except ‘flight’, ‘flight response’ and ‘nonphotochemical quenching’) were extracted together with their counts and summarised for polar day and polar night samples.

## RESULTS

### Seasonality

Our study spanned over 13 months and included two polar nights (three and two samples respectively), one polar day (five samples) and one sample from September coinciding with the transition period between polar day and polar night. Environmental parameters showed a seasonal pattern (Table 1, Supplement 2). This is a representative trend for the IsA time series station that displays a yearly recurrent pattern (Chitkara *et al*., unpublished data). Photosynthetically active radiation (PAR) at 25 m depth was detectable only between April and September 2012. Within this period the highest values were measured in April and beginning of May 2012, followed by the lowest detected values at the end of May and June. Hydrography of arctic fjords can be influenced by water masses originating from distinct sources and thus displaying different physiochemical properties categorised based on temperature and salinity [56], [57]. Locally formed cold water (< 1° C; LW) was present in the first half of the year (December 2011 to May 2012) with warmer intermediate water (>1° C; IW) influenced by land runoff and oxygen-rich Atlantic water dominated in the second half (Jun 2012 to Jan 2013). The coldest temperature was in January 2012 and the warmest in September 2012. Overall, nutrient concentrations (nitrate/nitrite, phosphate, and silicates) were heavily depleted from the onset of spring bloom until the end of polar day (from May to August; Table 1). Silicates, however, started to be depleted already in April (Table 1). Chlorophyll *a* was detectable throughout the year with a peak value in May and a second smaller peak in August. In all samples except those collected in May, most of the chlorophyll *a* was present in the small phytoplankton fraction (< 10 μm). Detailed descriptions of the IsA system, based on enhanced frequency of sampling can be found in [10], [12], [58], [59].

**Table 1.**
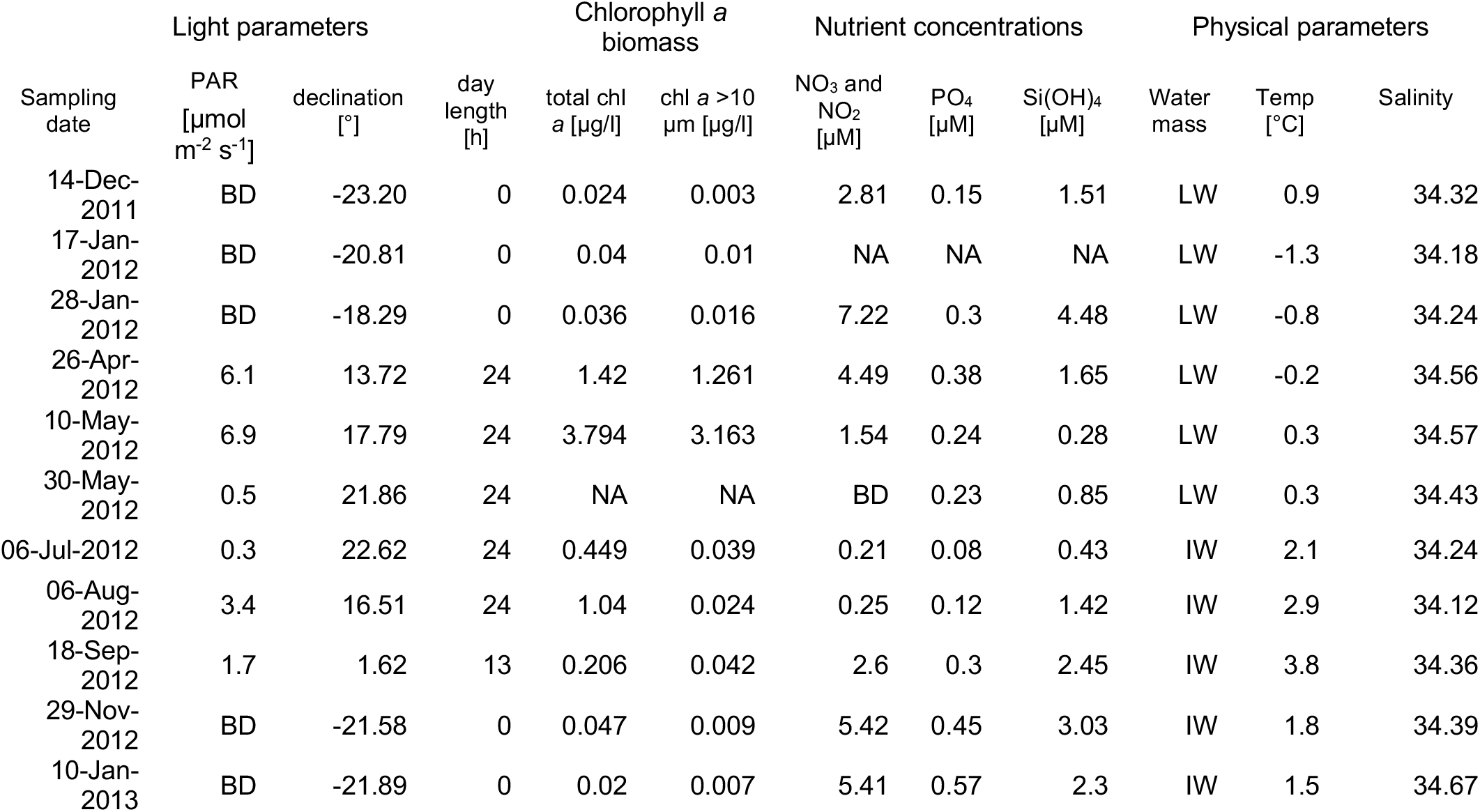
Environmental parameters corresponding to each metatranscriptome sampled at 25 m depth. PAR (photosynthetically active radiation) was measured as close to local noon as possible; declination was calculated for local noon, and day length refers to the number of hours when the sun is above the horizon. Chlorophyll *a* biomass is reported for 2 size fractions: total (filtered on GF/F glass microfiber filters (Whatman, England) and > 10 μm (filtered on Isopore membrane polycarbonate filters (Millipore, USA)). Water masses: LW – local water and IW – intermediate water. Other abbreviations: BD – below detection, NA – not available. The data were originally published in [56], [57].

### Seasonal transcript diversity

The diversity and evenness of transcripts was higher during polar night (n=5) than during polar day (n=5) (Figure 3). The mean number of transcripts collected during polar day was similar to the value obtained in September, during a mixed light regime (μ_PD_=1 178 988, σ_PD_=273 108 and 1 272 116, respectively), whereas average transcript diversity during polar night was 2.7 times higher. However, the January 2012 sample outlied significantly from the other polar night metatranscriptomes, containing 1.6 million transcript isoforms, a similar value to samples from polar day and September. Both the September 2012 and January 2012 samples that had low numbers of transcripts also had significantly lower depth of sequencing than the other samples (Supplement 1).

**Figure 3.**
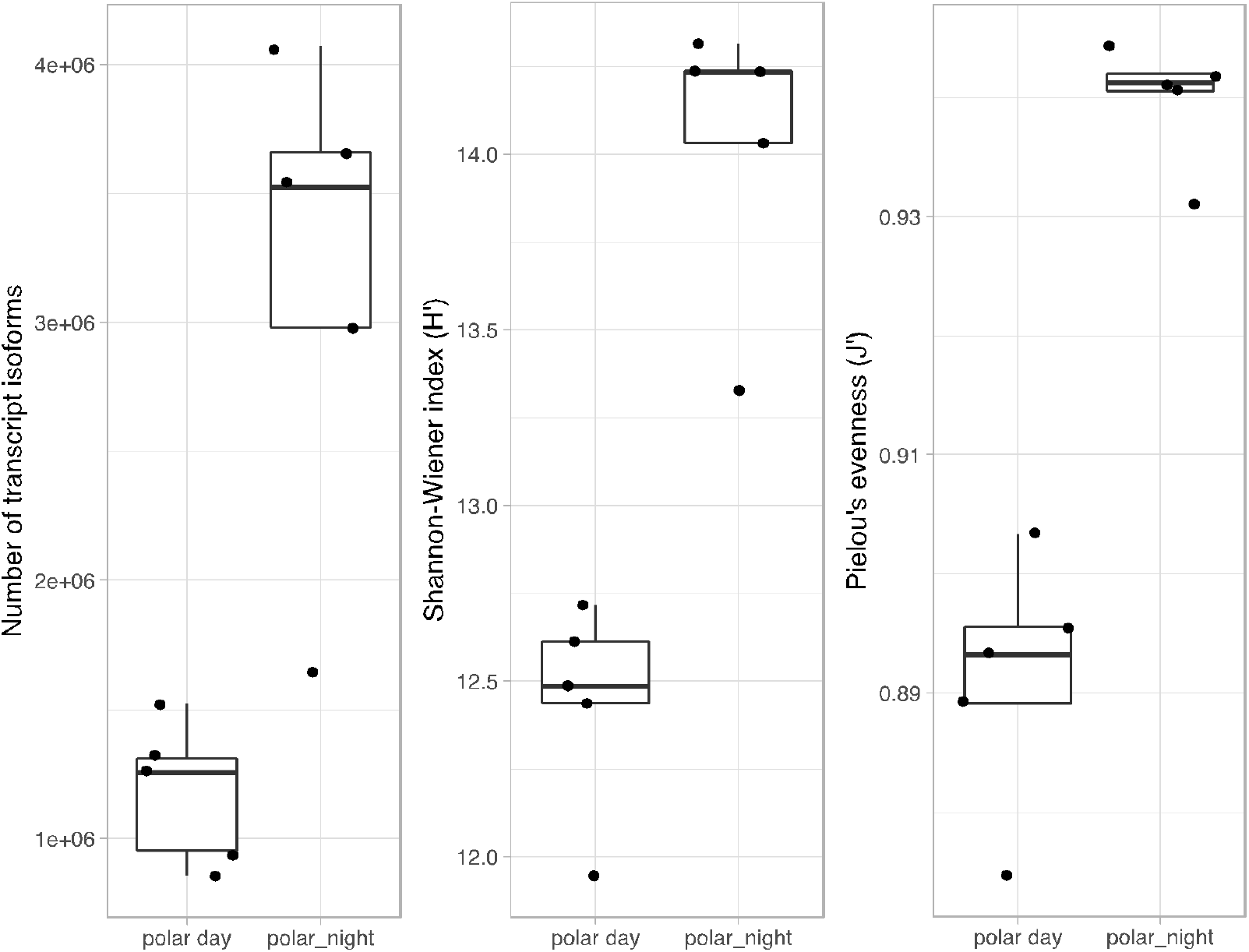
Diversity of transcript isoforms per sample during polar day (n=5) and polar night (n=5). The September sample was excluded.

We found a clear difference between metatranscriptomes from polar day and polar night with the September sample clustering with the polar night samples with high support (>99% of both unbiased and bootstrap probability; Figure 4). The polar day samples formed distinct subclusters (Figure 4). The core dataset containing almost 70 thousand of the most abundant transcripts showed similar or identical clustering, indicating that the pattern was not altered by the high contribution of rare transcripts (Figure 2 and 5). Further functional descriptions were therefore conducted using the core dataset.

**Figure 4.**
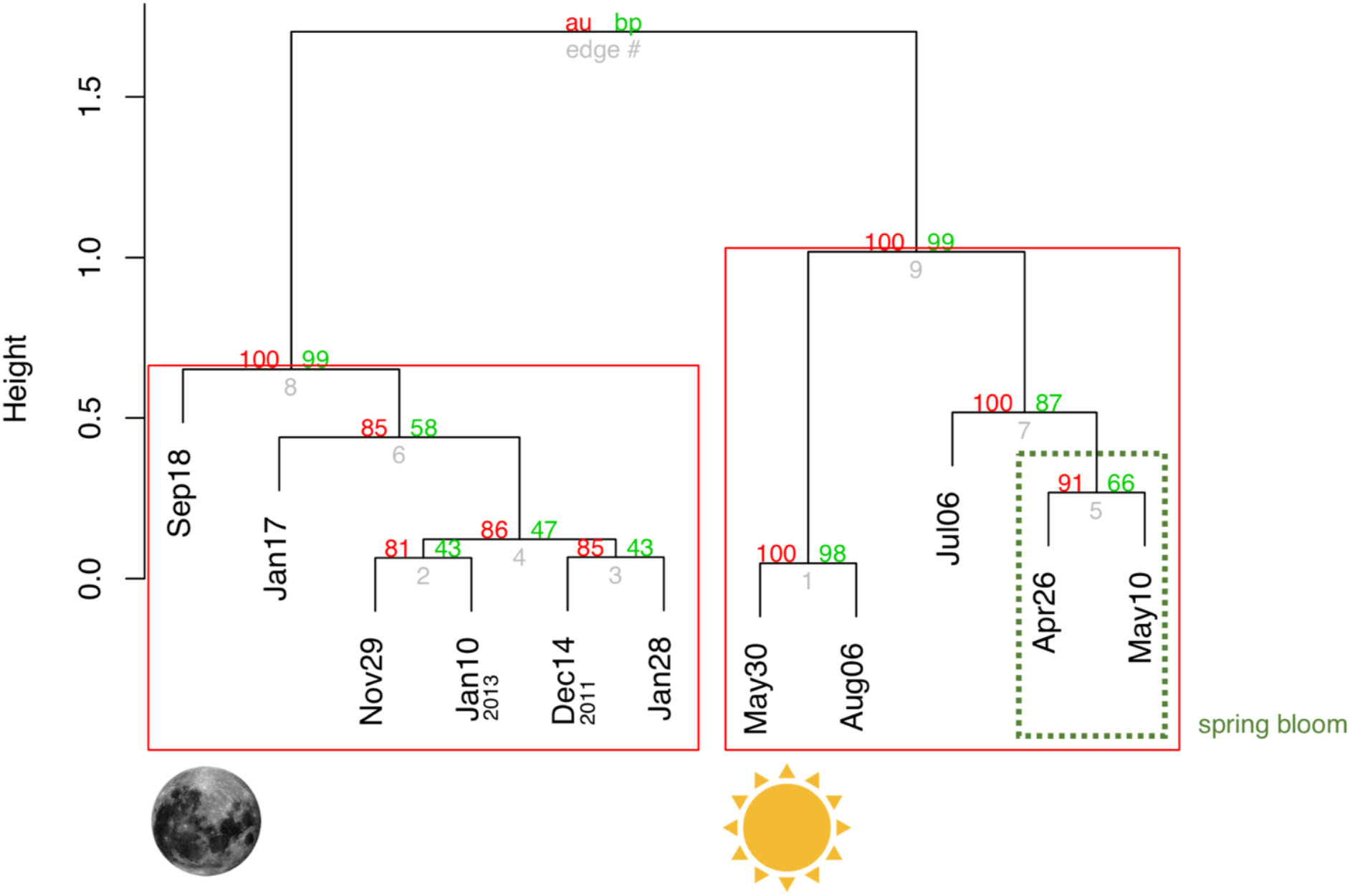
Grouping of the samples according to similarity in their transcript composition based on the core dataset. Approximately unbiased (au) and bootstrap probability (bp) values strongly support the clustering (au and bp > 80). Note that two main highly supported groups are delineated according to the light regime: polar day and times of the year with night present, i.e., polar day and September. The polar day cluster was divided into two groups with strong support (au and bp > 99).

**Figure 5.**
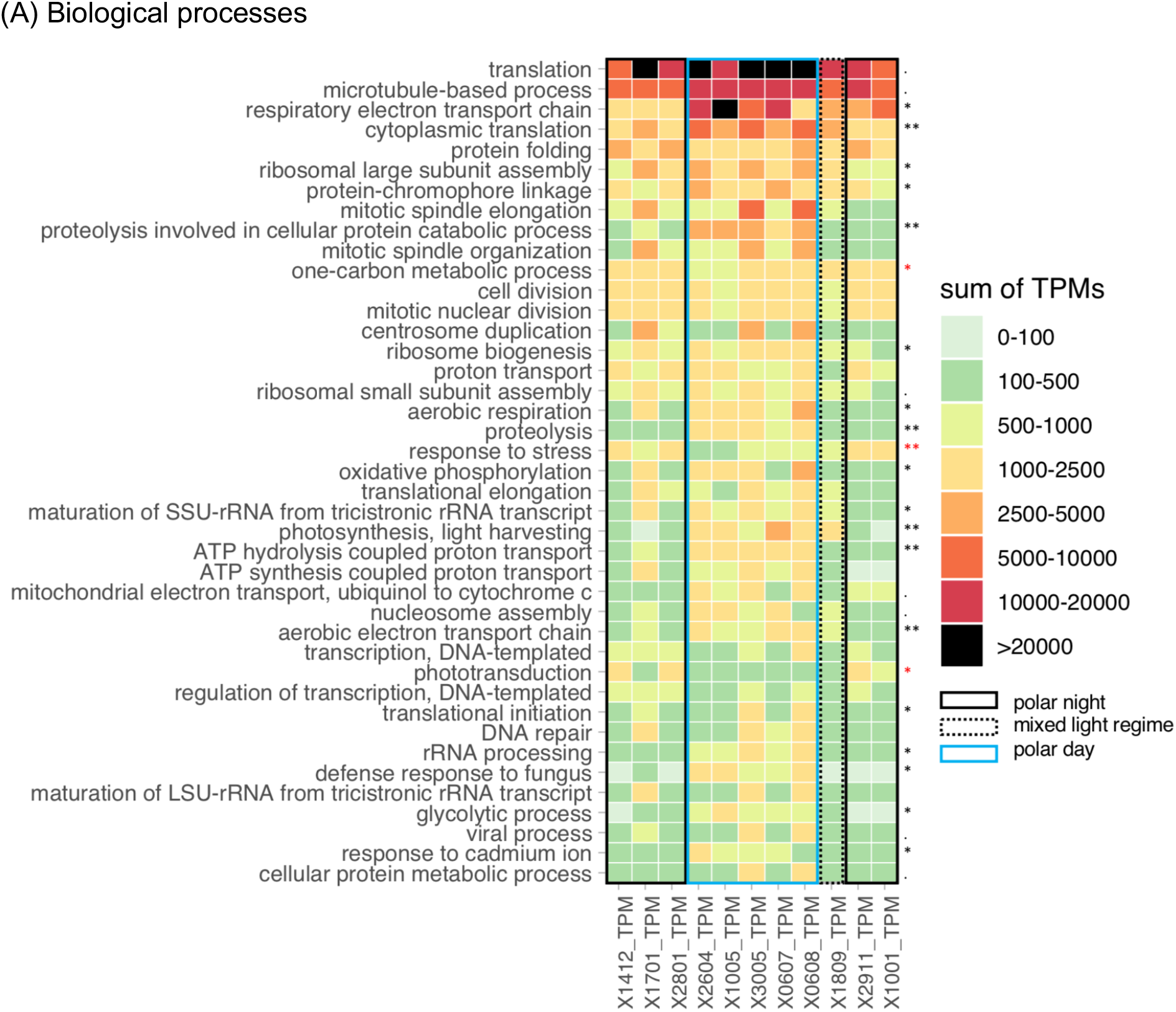
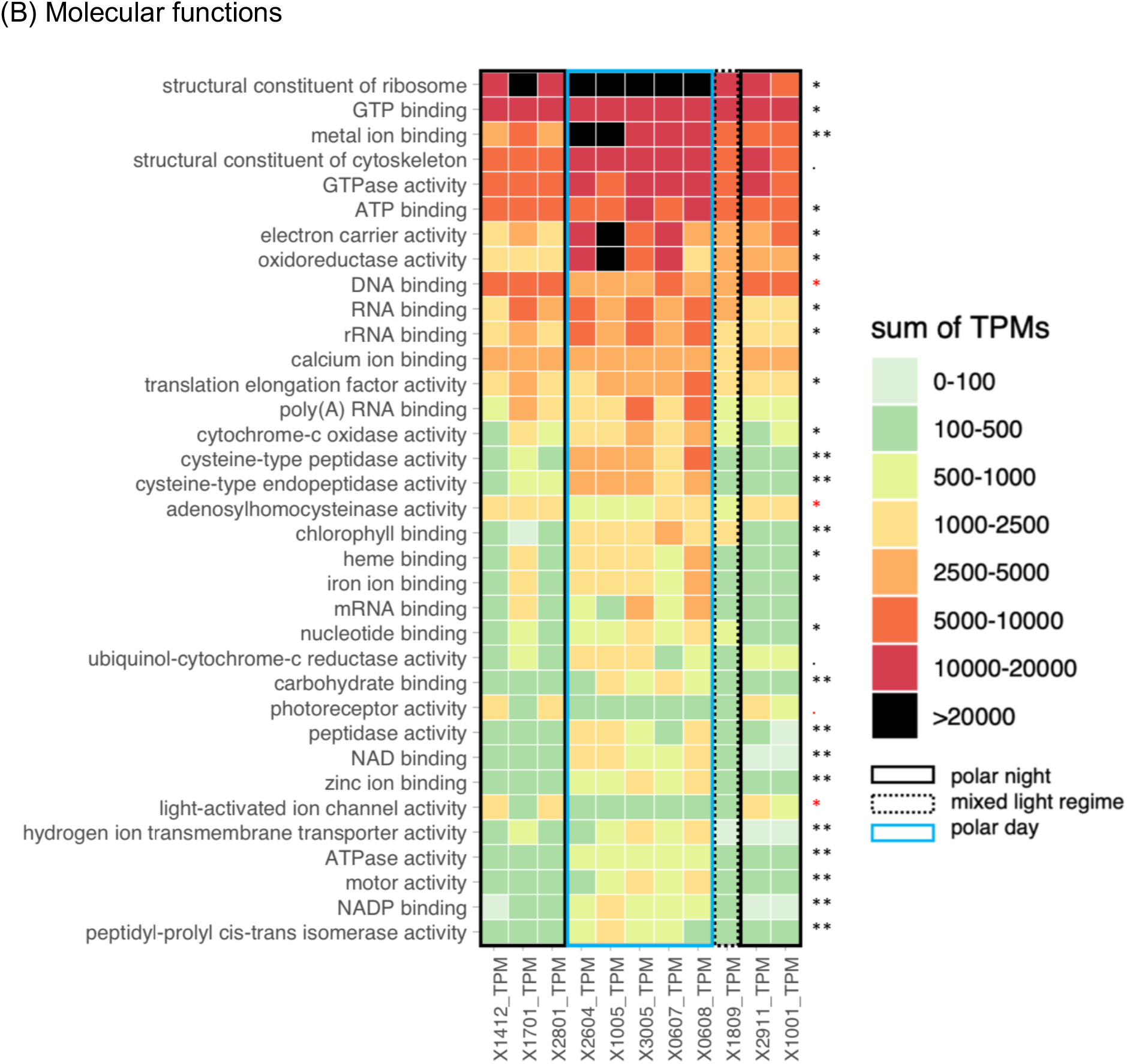
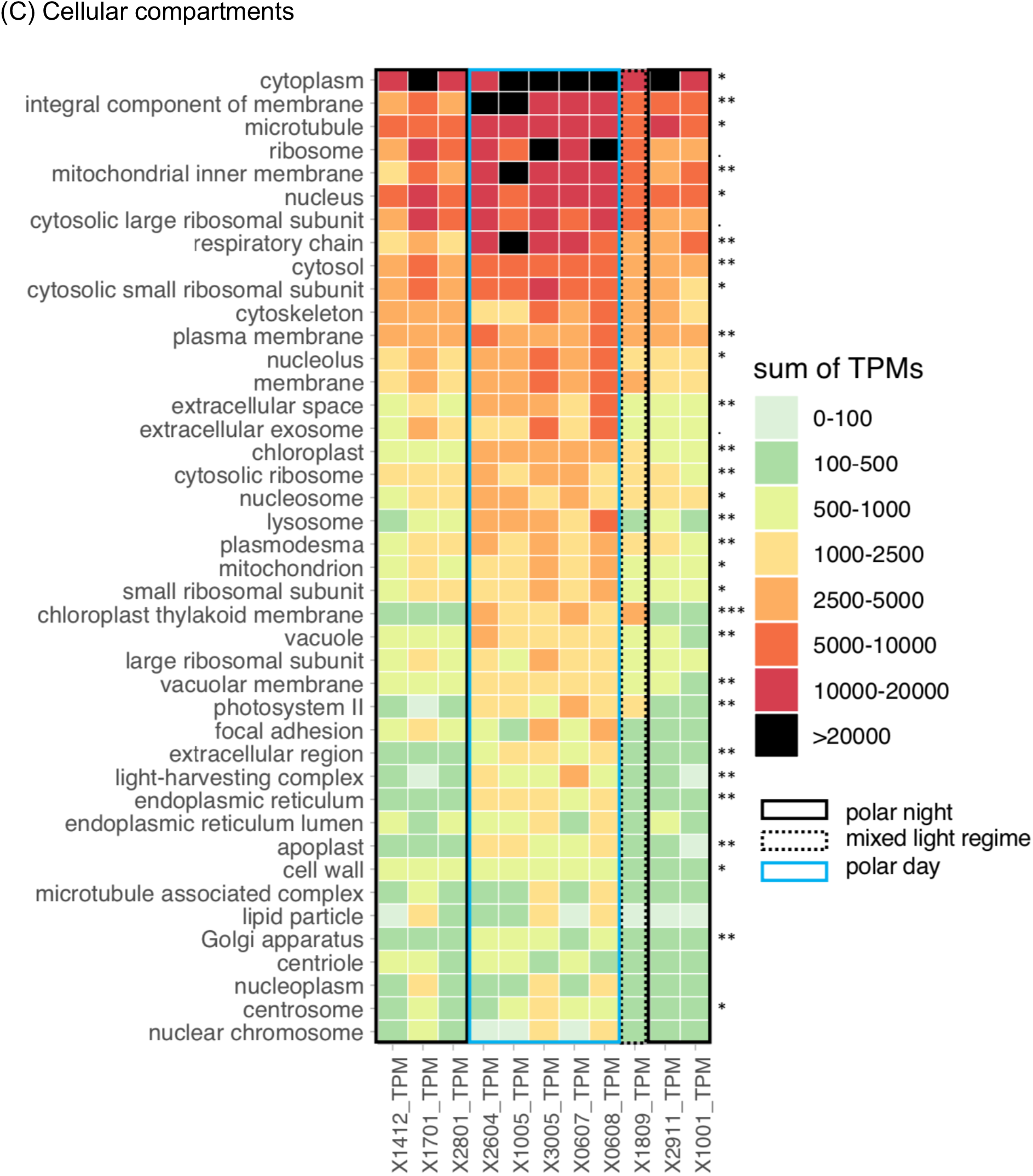
The most abundant GO terms within the core dataset corresponding to biological processes (A), molecular functions (B) and cellular compartments (C) featuring > 5000 TPM for each GO term. Asterisks indicate functions that differed between polar day (red asterisk) and polar night (black asterisk) using *simper* function (September was excluded from the Simper analysis). Significance codes: 0 ‘***’ 0.001 ‘**’ 0.01 ‘*’ 0.05 ‘.’ 0.1 ‘ ‘ 1.

We identified the transcripts with the strongest contribution to the differences between the main clusters (Figure 4). Ten of the transcripts contributing to the difference between polar night and polar day were also the most abundant transcripts in our dataset. Only the most abundant transcript out of these ten got a functional annotation and was classified as cytochrome b (Supplement 3).

### Taxonomic composition

The ratio of reads that could be assigned to taxonomic groups was similar throughout the year, ranging from 33 to 42% of all reads in each metatranscriptome. This left the majority of reads without a taxonomic annotation (58-67%). The proportion of taxonomically unannotated reads was independent of light regime and number of sequences per sample. The most represented supergroup in each sample was Alveolata, predominantly Dinophyceae and Ciliophora (Figure 8). Dinophyceae dominated in metatranscriptomes from polar night (32% on 17^th^ January 2012 up to 49% on 14^th^ December 2011) and September (33%), while ciliates were more abundant during polar day (18-34% versus 8-10% in polar night). Many taxonomic groups had low relative transcript abundance throughout the year, never exceeding 2% of the taxonomically assigned reads (Apusozoa, Bigyra, Cercozoa, Chromerida, Euglenozoa, Fornicata, Fungi, Glaucocystophyceae, Heterolobosea, Parabasalia, Perkinsea, Pseudofungi and Rhodophyta).

### Activity of expressed genes in a seasonal perspective

Annotation of the core dataset gene catalogue against the GO database resulted in 24,643 transcripts with at least one annotation (36% of core dataset). Environmental variables fitted into biological processes (GO category) dissimilarity matrix confirmed the importance of light as a structuring parameter (i.e., day length (R^2^_GNMDS_=0.88, p=0.019), declination (R^2^_GNMDS_=0.85, p=0.025) and PAR (R^2^_GNMDS_=0.54, p=0.082). On the other hand, the analysis did not support water mass (R^2^_GNMDS_=0.04, p=0.974) and temperature (R^2^_GNMDS_=0.20, p=0.475) as important explanatory variables in structuring biological processes.

The most abundant GO terms within biological processes belonged to housekeeping genes encoding proteins involved in translation, microtubule-based process, respiratory electron transport chain or protein folding etc. (Figure 5A). The majority of the most abundant biological processes were overrepresented in polar day samples, such as respiratory electron transport chain or cytoplasmic translation (Figure 5A). Some of the GO terms were more uniformly distributed throughout the year, such as cell or mitotic nuclear division (Figure 5A). Finally, a few of the most abundant GO terms were overrepresented during polar night. This was the case for one-carbon metabolic processes (mean number of TPM in polar night samples, μ_PN_=1974, μ_PD_=1134 in polar day samples, p=0.03), response to stress (μ_PN_=1482 in polar night, μ_PD_=498 in polar day, p=0.01) and phototransduction (μ_PN_=936 in polar night, μ_PD_=323 in polar day, p=0.03). The majority of transcripts within one-carbon metabolic processes mapped to adenosylhomocysteinase and S-adenosylmethionine synthase. The latter catalyses hydrolysis of L-methionine into S-adenosyl-L-methionine which is an essential source of different chemical groups, e.g. methyl groups used for epigenetic modifications including DNA methylation [60], [61]. Whereas adenosylhomocysteinase catalyses one of the next reactions in methionine metabolism: hydrolysis of S-adenosyl-L-homocysteinase to adenosine and L-homocysteine [62] and has been connected to silicon [63] and vitamin starvation in diatoms [64]. All transcripts in response to stress mapped to chaperone proteins, most (451 out of 456) mapped to different types of heat shock proteins, especially HSP90 (406 transcript isoforms).

Almost all light-dependent biological processes were relatively more abundant in polar day samples (Figure 7). This was especially true for GO terms connected to photosynthesis. However, most of the categories were also present during polar night albeit in low numbers. Three of the terms were more abundant in polar night, such as eye photoreceptor cell development, phototaxis and especially phototransduction. Phototransduction contained 208 transcripts mapping to green- and blue-light absorbing proteorhodopsins.

Most transcripts contributing to less abundant GO terms, but overrepresented during polar night (Figure 6), mapped to multipurpose proteins, mainly chaperones (HSP72 and HSP71 in protein refolding, HSP72 in negative regulation of cellular response to heat or response to virus). Phagocytosis and response to other organism categories consisted mostly of transcripts assigned to calreticulin, a multipurpose protein acting as calcium-level regulator and chaperone in endoplasmic reticulum [65]. Pathogenesis contained mostly tripeptidyl-peptidase transcripts and acidic proteases probably involved in virulence response [66]. Response to cycloheximide, a naturally occurring fungicide, contained transcripts mapping to 60S ribosomal protein L44.

**Figure 6.**
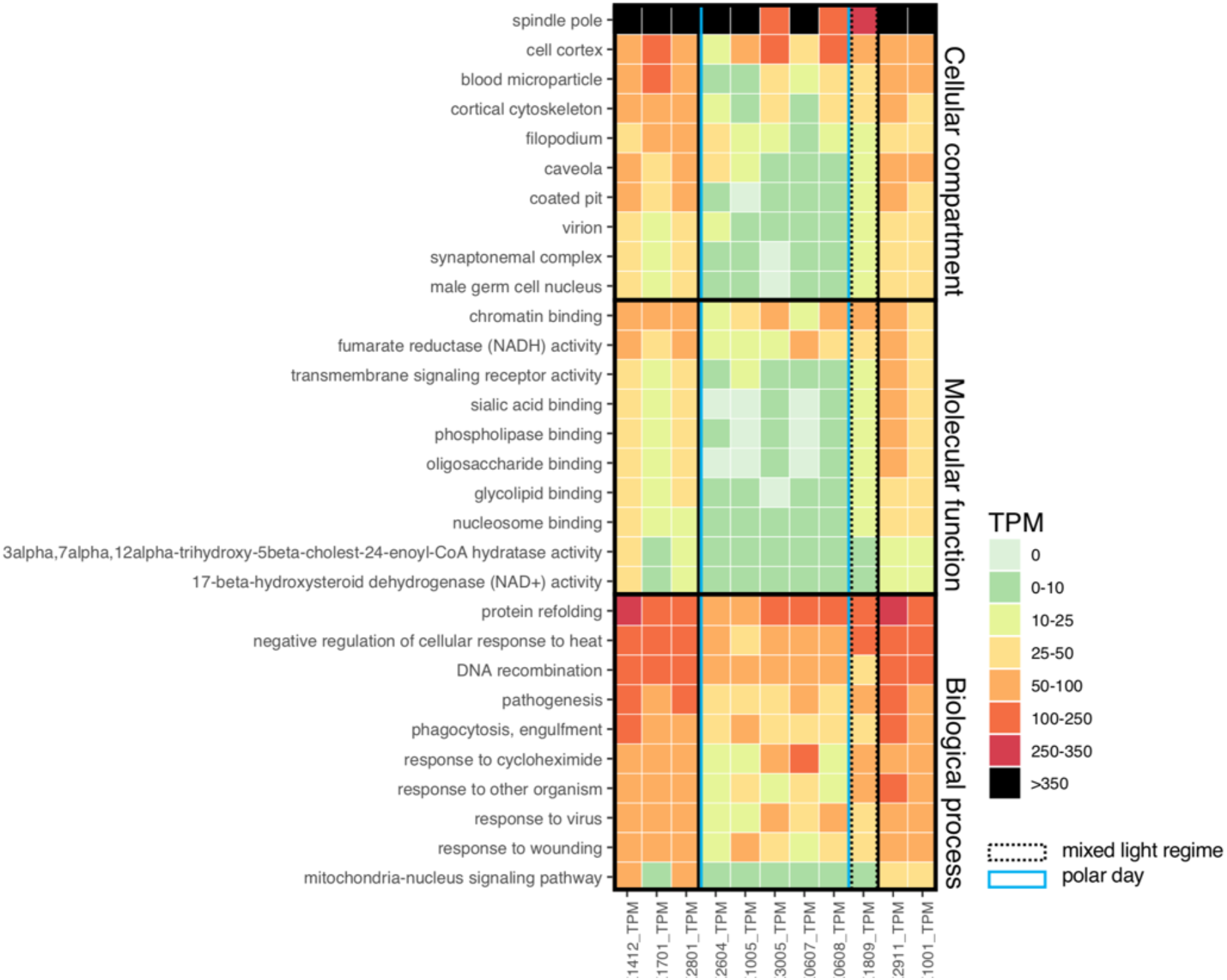
Top 10 less abundant GO terms (sum of TPM in all metatranscriptomes < 5000) with mean number of transcripts higher during polar night than during polar day (p > 0.05).

Most GO terms within molecular functions were overrepresented in polar day (Figure 5B). Analyses indicated light, but not water masses, to be an important structuring factor of the most abundant molecular functions of the community (R2_GNMDS_=0.795, p=0.005 versus R2_GNMDS_=0.017, p=0.897, respectively). Only DNA binding (μ_PN_=6766 and μ_PD_=4714, p=0.024), adenosylhomocysteinase activity (μ_PN_=1585 and μ_PD_=889, p=0.017), photoreceptor (μ_PN_=936 and μ_PD_=315, p=0.055) and light-activated ion channel activity (μ_PN_=883 and μ_PD_=247, p=0.025) were overrepresented in polar night. DNA binding is a broad category of gene products that have been identified as reacting selectively in a non-covalent manner with DNA. We identified 1651 transcripts containing mostly major basic nuclear proteins, histones, cold shock proteins etc. Light-activated ion channel and photoreceptor consisted mostly of identified proteorhodopsins; additionally, photoreceptor contained also transcripts mapping to centrins. Centrins are calcium-binding proteins involved in centrosome and microtubule functioning [67], as well as regulation of signalling and molecular translocation [68]. Among less abundant molecular functions overrepresented during polar night, we found that the transcripts mapped mostly to multipurpose proteins, similarly to biological processes. Chromatin binding consisted of diverse proteins, with the majority of transcripts mapping to 60S ribosomal protein L29. Fumarate reductase (NADH) activity consisted of transcripts mapping to an enzyme that catalyses reversible anaerobic reduction of succinate to fumarate, generating NADH and protons [69]. Sialic acid, phospholipase and oligosaccharide binding contained transcripts mapping mainly to e-selectin, a protein involved in an inflammatory response that changes properties of the cell surface [70]. Mapping to heat shock-related 70kDa proteins was found in glycolipid binding, whereas nucleolin in nucleosome binding. Nucleolins are also plurifunctional proteins that play important roles in viral infections [71]. 17-beta-hydroxysteroid dehydrogenase (NAD+) activity and 3alpha,7alpha,12alpha-trihydroxy-5beta-cholest-24-enoyl-CoA hydratase activity contained the same transcript isoforms that mapped to peroxisomal multifunctional enzymes taking part in β-oxidation of lipids [72] but could also be necessary in fungal pathogenesis [73].

The strongest differences in transcripts within the three GO categories between polar day and polar night was found in cellular compartment (R_ANOSIM_=0.928, p=0.01); with less pronounced differences in biological processes (R_ANOSIM_=0.792, p=0.008) and molecular functions: (R_ANOSIM_=0.892, p=0.009). At the same time among the most abundant transcripts divided according to the respective cellular compartments, we did not find any category that would be overrepresented during polar night (Figure 5C). Less abundant GO terms pointed out compartments overexpressed during polar night that are connected to cytoskeleton (spindle pole, cell cortex, cortical skeleton, filopodium) or cell membrane (coated pit, cell cortex, caveola; Figure 6). It should be noted that the names of some categories are misleading for microbial eukaryotes, such as male germ cell nucleus (containing the peroxisomal multifunctional enzymes mentioned in molecular functions) or blood microparticle (containing mostly actins or signalling molecules). The spindle pole term consisted of transcripts that mapped to centrins, mentioned in the photoreceptor term in molecular functions. Cell cortex region lies beneath the cell membrane and contained transcripts that mapped to myosins, 14-3-3-like proteins, profilins and other cytoskeleton related proteins.

### Transcript novelty

Levels of functional annotation were overall low, regardless of the database used. Mapping to UniProt (with blastp), Pfam, TmHMM, GO (based on Pfam) resulted in <10% of transcript annotation, while eggNOG and KEGG gave 10-20% successful annotation. Only UniProt (with blastx) and GO (with blastp) annotated 38% and 36% of transcripts, respectively. However, mapping our assembled transcripts to the *Tara* Oceans datasets showed that most of our transcripts had hits, matching especially samples from the Arctic (Figure 9). Up to 75% of our transcript isoforms mapped to the surface samples (station 196, north of Alaska), up to 78% mapped to the deep chlorophyll maximum layer (station 173, northeast of Novaya Zemlya), and up to 74% to the mesopelagic zone (station 201 in west part of Baffin Bay). The mean proportion of transcripts mapping to surface samples from *Tara* Oceans stations located north of 60°N was much higher than for stations in the temperate and tropical regions (μ_↑60N_=64%, σ_↑60N_=9% and μ_↓60N_=21%, σ_↓60N_=8%, respectively). This was also true for samples from the deep chlorophyll maximum depth (μ_↑60N_=69%, σ_↑60N_=12% and μ_↓60N_=23%, σ_↓60N_=9%, respectively) and mesopelagic depths 67% (σ_↑60N_=6%).

**Figure 7.**
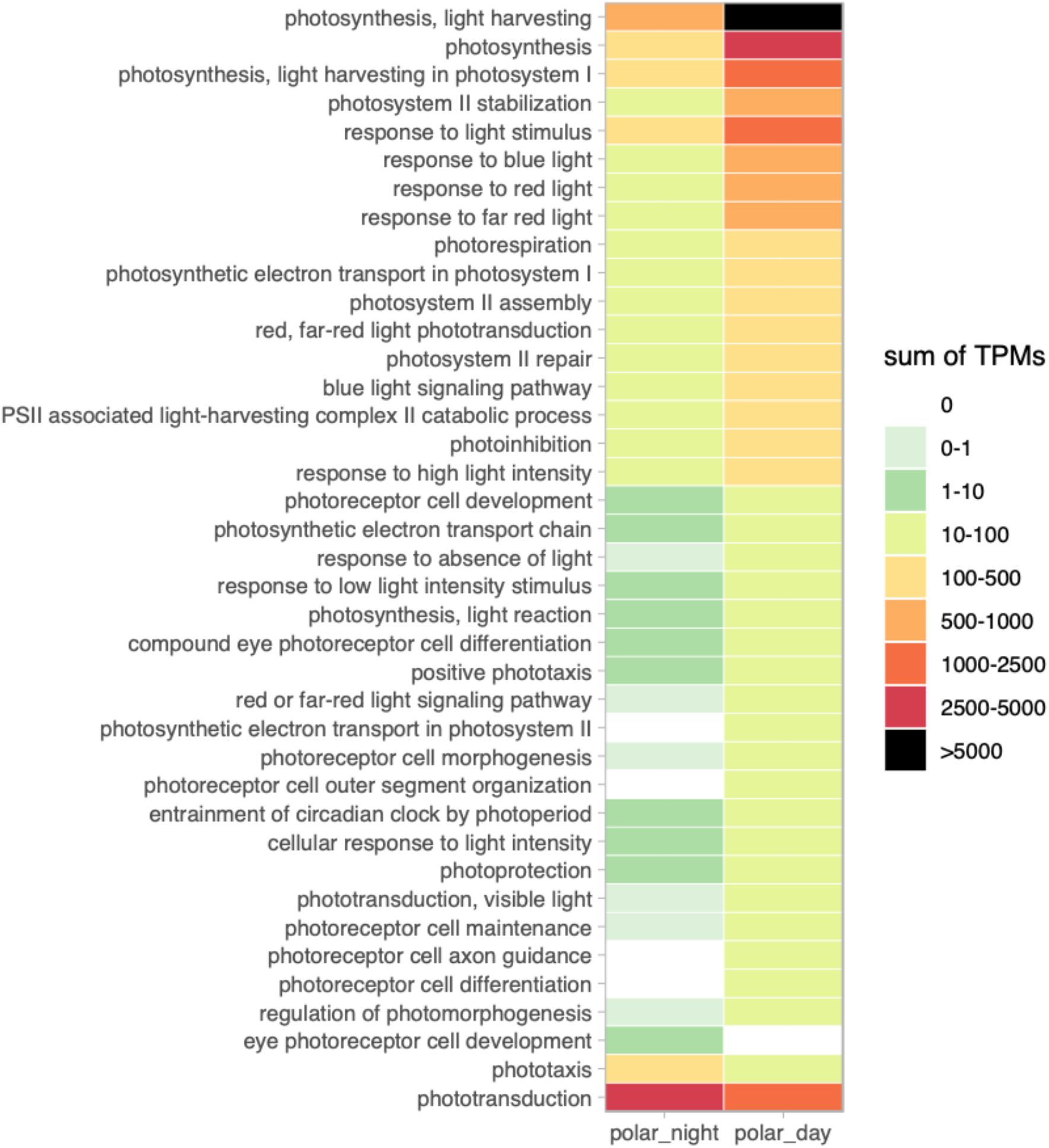
Seasonal abundances of transcripts associated with light-dependent biological processes (GO terms), shown as sum of transcripts per million (TPM) for samples from polar night (n=5) and polar day (n=5).

**Figure 8.**
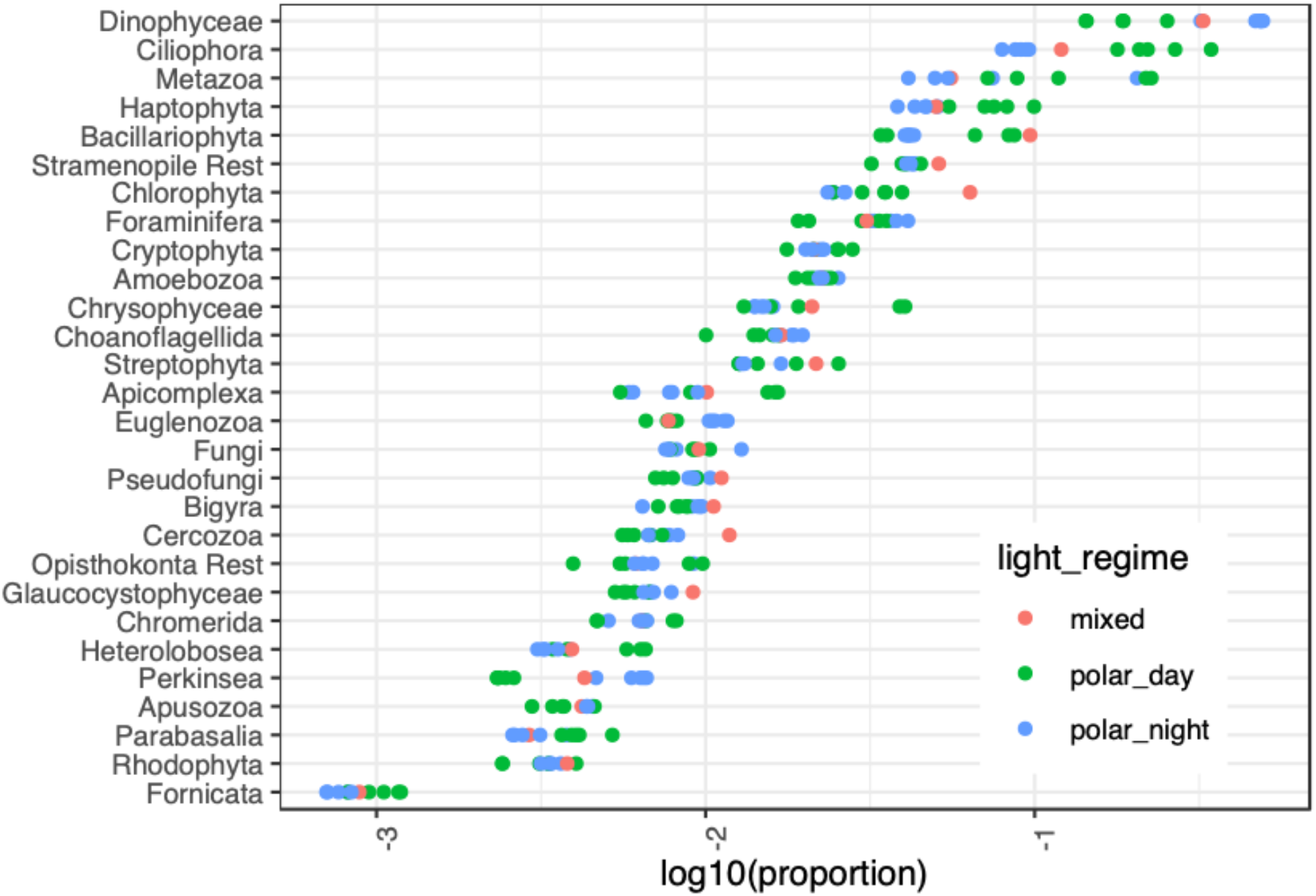
Taxonomic assignment shown as the proportion of clean reads assigned to a taxonomic group with Taxmapper. Each dot represents the proportion of reads in one sample.

**Figure 9.**
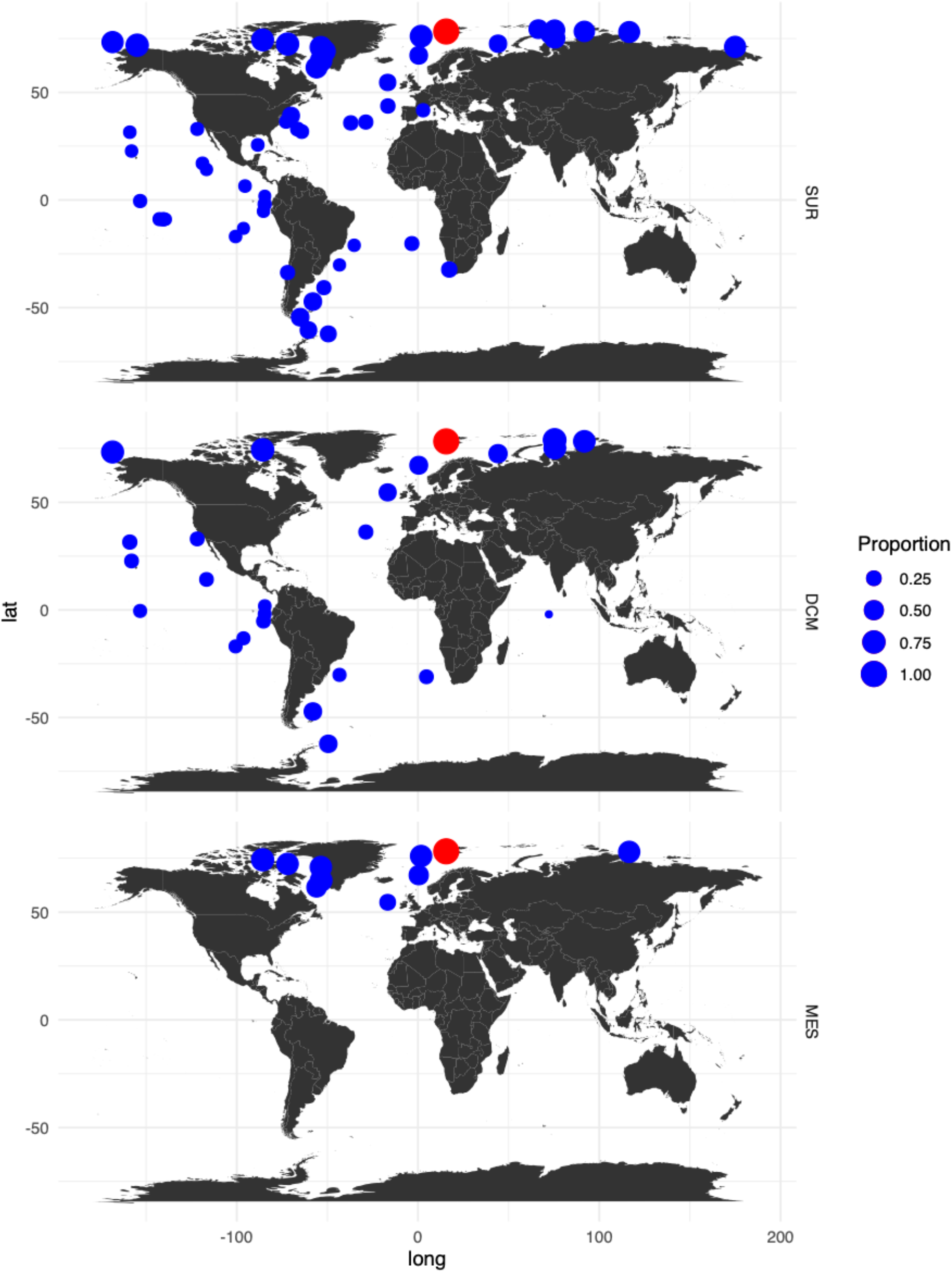
Proportion of transcripts isoforms from IsA core dataset (red circle) and proportion of their matches to data from *Tara* Oceans stations (blue circles). *Tara* Oceans data from 0.8-2000 μm plankton size fraction collected at different depths (SUR – surface waters, DCM – deep chlorophyll maximum and MES – mesopelagic waters).

## DISCUSSION

Climate change is already influencing Arctic marine ecosystems [74] and different scenarios for the development of Arctic marine ecosystems have been suggested. However, predicting its influence on polar ecosystems is challenging without a deep understanding of both the structure and function of its components [75]. Thus, the responses of microbial communities to these changes cannot be predicted without understanding which biological and molecular activities are taking place and how they impact biogeochemical cycles. Differences in gene expression could change the outcome of trophic interactions in an ecosystem, potentially altering the energy and nutrient flow to higher trophic levels [26], [75]. In this study we went beyond reporting detected species or its molecular proxies by examining community-level molecular engagement in biological activities. Our study offers a first description of the key processes performed by the microbial eukaryotic community over seasons in the Arctic fjord.

The strong seasonality at high latitudes affects microbial eukaryotes by influencing cell counts, biomass distribution, community composition, dominating carbon acquisition mode and various biodiversity measures [10], [12], [76]. Therefore, seasonal gradients profoundly affect the overall pool of present genes and their products, i.e., gene transcripts or proteins. Higher richness (number of operational taxonomic units (OTUs)) of marine protists during polar night compared to polar day was described independently in distant parts of the arctic marine waters (e.g., [10], [77]). Same patterns were shown for other arctic marine microorganisms, such as bacteria and archaea [78]. This likely panarctic phenomenon could originate from physically driven mixing throughout the water column which could enable the detection of diverse microorganisms at various atypical water depths during polar night [e.g., 14]. In this sense, mixing increases species diversity at different water depths. Temperature and salinity profiles during polar night are uniform throughout the water column at IsA [10] and thus could support this explanation. We did not find unequivocal evidence for increased functional diversity in microbial eukaryotes’ transcript, i.e., expression of a wider array of genes needed for survival (data not shown). In line with previously published evidence for higher richness during polar night, we showed that diversity and evenness of transcripts were also higher during polar night. The proportions of transcripts belonging to predominantly photosynthetic protists such as diatoms, haptophytes and chlorophytes, were consistently lower during polar night, confirming lower representation in the community and perhaps also lower overall activity of organisms in these groups [10], [14]. However, despite high diversity of OTUs and transcripts, cell counts, and therefore biomass of protists remained low throughout polar night [12], [79]–[81].

During polar night the contribution of single species to the overall low pool of biomass is more even than at any other times of the year, especially spring bloom [10], [12]. This includes crucial primary producers, such as *Micromonas polaris*, which were encountered as active at different depths of Arctic marine habitats during polar night [14]. Persistence of low levels of light-dependent biological processes in primary producers during polar night is likely due to the persistence and perhaps even maintenance of a functional photosynthetic apparatus kept ready to be activated once the light comes back [13], [30]. Therefore an overrepresentation of eukaryotic proteorhodopsins during polar night was rather unexpected, as bacterial proteorhodopsins are known to contribute to an alternative pathway to photosynthesis being the main contributors to harnessing solar energy in the ocean [82]. It is not clear what is their function in microbial eukaryotes, such as dinophytes [83]. However, recently, it was suggested that they are involved in G protein-coupled receptor-based signalling in Dinophyta [84].

Gene expression is likely to be more strictly controlled in many organisms during polar night due to overall lower availability of energy in the ecosystem [4]. In our dataset we found several GO functions that might hint to expression of genes that are involved in energy conservation. An increased expression of histones or major binding nuclear proteins or similar genes could serve as a way to control gene expression by binding and thus preventing DNA from being transcribed [85]. On the other hand, it may also point towards cellular division and the need to produce new histones for new cells [86]. GO term classification of transcripts overrepresented during polar night covers mostly categories such as response to stress, cellular signalling, modifications in cytoskeleton, pathogenesis, etc., through proteins that are known to be multifunctional. Multifunctionality might be an important strategy for efficient use of resources that could limit some groups of organisms during polar night. Other functions overrepresented during polar night involve adenosylhomocysteinase that could play an important role in increasing lifespan of microbial eukaryotes by controlling the concentration of methionine [87]. In general, biochemical reactions involved in methionine degradation are the main source of methyl groups used in gene silencing by DNA methylation which could possibly be another argument for strict control of gene expression during polar night [87]. Overrepresentation of different types of chaperon and heat shock protein transcripts during polar night may be connected to high demand of energy conservation by assuring correct assembly, maintenance and stability of proteins’ structures within the cell [88], [89]. Moreover, heat shock proteins could influence increased cell survival by several mechanisms able to attenuate apoptosis [90].

Among the most expressed transcripts in our study were a few functionally annotated sequences reaching up to 38% of the total number of transcripts which coincides with similar numbers of taxonomically annotated reads in our study. Metatranscriptomic studies often report low levels of functional annotations (down to 19%) that might be a result of various factors, such as the complexity of the studied environment [91], available reference databases [49], choice of algorithms, bioinformatic tools and parameters used for data analysis [92], etc.

To date, the most extensive marine global survey examining expressed eukaryotic genes based on *Tara* Oceans 2009-2012 reported 51.2% unannotated clusters of expressed genes [49]. Although the overall rates of annotation in our study were low, the data mapped successfully against the *Tara* Oceans dataset (including *Tara* Oceans Polar Circle campaign in 2013) by matching to 78% of transcript isoforms, specifically in polar regions (Figure 9). Therefore, we conclude that *de novo* assembled transcripts in our bioinformatic pipeline were robust and contained 22% of novel genes that are less likely to be found at lower latitudes (Figure 9). We hypothesize that the proportion of successfully mapping transcripts in our study would have been higher if the *Tara* Oceans campaign in the Arctic was extended beyond June-October to collect samples during polar night.

The high proportions of transcripts mapping to *Tara* Oceans’ metatranscriptomes from the Arctic suggest a distinct genetic makeup of microbial eukaryotes in this part of the world. Perhaps the different genetic makeup of eukaryotic communities in high latitudes could reflect necessary adaptations to Arctic seasonality that are not present in potential invasive microbial eukaryotes moving northwards with progression of climate warming. Recent studies on marine biogeography of DNA viruses revealed their biodiversity hotspot in the Arctic Ocean [93]. Since viruses and mobile elements carried by them are known as powerful agents of evolution in all living cells [94], they could potentially contribute to increased diversification of genes in microbial communities in the Arctic [95], resulting in an observed low similarity to other parts of the ocean. This topic requires a separate line of research, and regardless of possible links between the two groups, the distinct genetic makeup of microbial eukaryotes stands in need for more exploration.

Polar night seems to work as a reset stage for Arctic marine environments, possibly enforcing shifts to heterotrophy in the absence of light and allowing protist survival as low biomass populations. Moreover, changing proportions of transcripts annotated to taxonomic groups as well as fluctuating abundances of functional categories point out that community-level metabolic state changes together with shifting community composition. The two polar nights in our study showed a striking similarity in taxonomic and functional composition of transcripts that might reflect a specific, recurrent impact of environmental filtering imposed by seasonal light regime and temperature. A long-term monitoring of taxonomic and transcriptional dynamics could evaluate to which extent other factors such as inflow of warmer water masses or arrival of species moving northwards, influence the strength of light regime filtering and development of future eukaryotic communities.

## Data availability

The data for this study have been deposited in the European Nucleotide Archive (ENA) at EMBL-EBI under accession number PRJEB48707 (https://www.ebi.ac.uk/ena/browser/view/PRJEB48707, access from 1^st^ December 2021). Raw data from *Tara* Oceans are available at EBI and GenBank under project IDs PRJEB9738 and PRJEB9739.

## Acknowledgments

The authors are thankful to all the people that contributed to the marine part of MicroFun project, collected and processed the samples. This research was funded by University Centre in Svalbard, as well as ConocoPhillips and Lundin Petroleum through The Northern Area Program and by the Norwegian Research Council, project number 230970. The cost of sequencing was partly covered through Jan Christensens Legat. This work was performed on the Abel Cluster, owned by the University of Oslo and the Norwegian Metacenter for High Performance Computing (NOTUR), and operated by the Department for Research Computing at USIT, the University of Oslo IT-department. http://www.hpc.uio.no/. The publication charges for this paper have been funded by a grant from the publication fund of UiT - The Arctic University of Norway. We are also deeply thankful to the *Tara* Oceans Consortium who provided open and freely available data.

## Author contributions according to CRediT – Contributor Roles Taxonomy

MW: Formal analysis, Writing - Original Draft, Writing - Review & Editing, Visualisation

AV: Formal analysis, Resources, Writing - Original Draft, Writing - Review & Editing

RL: Formal analysis, Resources, Writing - Review & Editing

EP: Formal analysis, Resources, Writing - Review & Editing

TMG: Resources, Writing - Review & Editing, Funding acquisition

## Competing interests

The authors declare no competing interests.

## Notes

### Competing Interest Statement

The authors have declared no competing interest.

