## Supplement for "Linking extreme seasonality and gene expression in arctic marine protists"

### Supplement 1

Pre-processing statistics of the sequenced samples.

**Supplement 2**

Principal component analysis of environmental data. 17 January 2012 and 30 May 2012 are not included in the figure, due to some missing values.

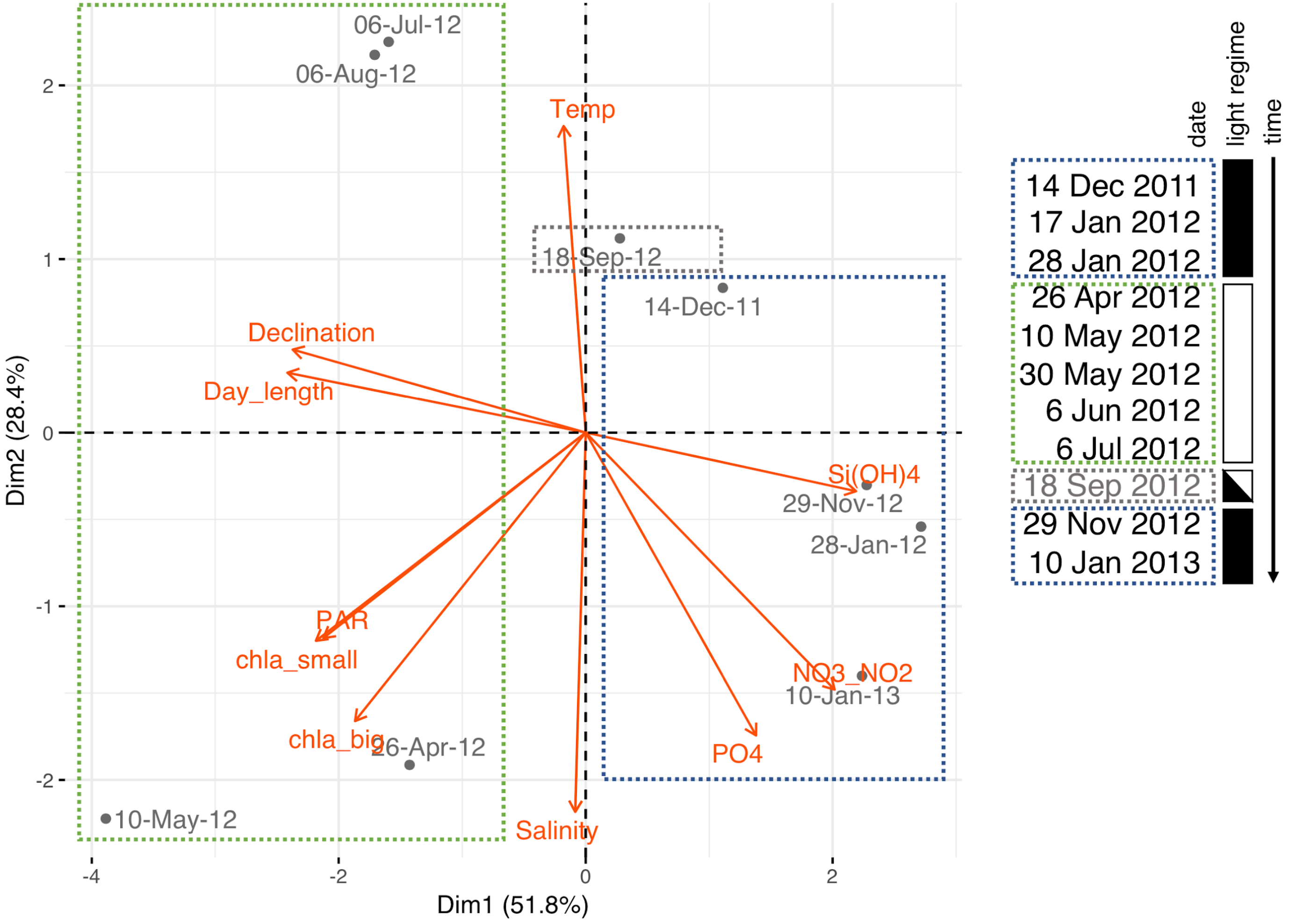

**Supplement 3**

Most transcript isoforms were characterized by low sum of relative abundance across samples: 8 transcripts with >10 000 TPM, 154 with >1000 TPM, 3483 with >100 TPM, 68 166 with > 10 TPM and 2 390 862 with >1TPM. Most of the least abundant transcripts were unannotated with the Gene Ontology (based on blast results from mapping transcript isoforms to UniProt).

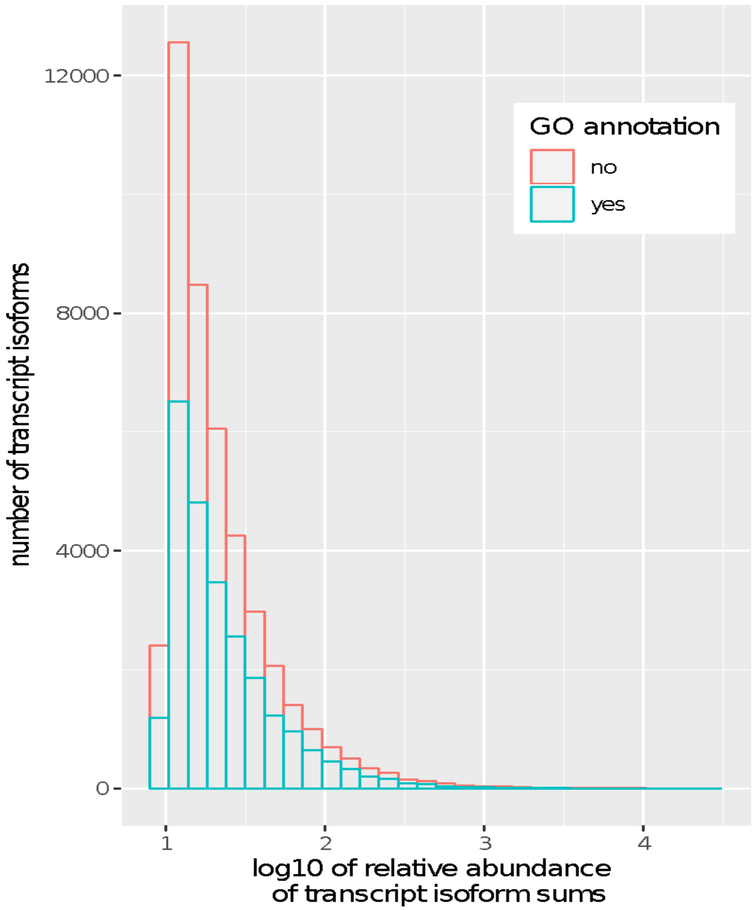

**Supplement 3**

Summary of results from SIMPER analysis of Bray-Curtis dissimilarity matrix of reduced dataset. These are the 10 top individual transcript isoforms contributing to differences between grouping factors (transcript in bold was the only one among the top 10 in section a, b, c that was functionally annotated by at least one database; yellow highlight – transcript isoforms present in a, b and c).

a. grouping factor: polar night vs. polar day (without September)

| | average | sd | ratio | ava | avb | cumulative sum |
|---|---|---|---|---|---|---|
| **TRINITY_DN3567435_c13_g1_i1** | **0.0091068** | **7.643e-03** | **1.1916** | **519.77** | **5236.75** | **0.01020** |
| TRINITY_DN3531545_c4_g1_i13 | 0.0057171 | 4.039e-03 | 1.4154 | 645.86 | 3522.82 | 0.01661 |
| TRINITY_DN3383058_c4_g1_i10 | 0.0049891 | 4.020e-03 | 1.2409 | 210.17 | 2793.45 | 0.02219 |
| TRINITY_DN3531545_c4_g1_i11 | 0.0044515 | 2.692e-03 | 1.6536 | 572.16 | 2830.47 | 0.02718 |
| TRINITY_DN3528005_c14_g2_i1 | 0.0033457 | 1.859e-03 | 1.7995 | 51.95 | 1762.08 | 0.03093 |
| TRINITY_DN3567090_c10_g1_i2 | 0.0032716 | 2.241e-03 | 1.4599 | 411.10 | 2057.50 | 0.03459 |
| TRINITY_DN3565854_c4_g3_i5 | 0.0032060 | 4.265e-03 | 0.7518 | 2.79 | 1660.42 | 0.03818 |
| TRINITY_DN3417822_c3_g2_i11 | 0.0031592 | 2.095e-03 | 1.5081 | 271.12 | 1859.83 | 0.04172 |
| TRINITY_DN3313835_c6_g2_i3 | 0.0027240 | 2.927e-03 | 0.9306 | 45.65 | 1396.98 | 0.04477 |
| TRINITY_DN3417822_c3_g2_i2 | 0.0024838 | 2.791e-03 | 0.8898 | 397.70 | 1458.13 | 0.04756 |

b. absence vs. presence of night

| | average | sd | ratio | ava | avb | cumulative sum |
|---|---|---|---|---|---|---|
| **TRINITY_DN3567435_c13_g1_i1** | **0.0089522** | **0.0075762** | **1.1816** | **635.64** | **5236.74** | **0.01014** |
| TRINITY_DN3531545_c4_g1_i13 | 0.0055438 | 0.0039526 | 1.4026 | 780.39 | 3522.82 | 0.01642 |
| TRINITY_DN3383058_c4_g1_i10 | 0.0048617 | 0.0040257 | 1.2077 | 279.26 | 2793.45 | 0.02193 |
| TRINITY_DN3531545_c4_g1_i11 | 0.0042184 | 0.0026293 | 1.6044 | 739.23 | 2830.47 | 0.02670 |
| TRINITY_DN3528005_c14_g2_i1 | 0.0033084 | 0.0018580 | 1.7807 | 74.10 | 1762.08 | 0.03045 |

| | | | | | | |
|---|---|---|---|---|---|---|
| TRINITY_DN3565854_c4_g3_i5 | 0.0032128 | 0.0042582 | 0.7545 | 2.32 | 166.42 | 0.03409 |
| TRINITY_DN3567090_c10_g1_i2 | 0.0031939 | 0.0022026 | 1.4501 | 469.78 | 2057.50 | 0.03771 |
| TRINITY_DN3417822_c3_g2_i11 | 0.0030590 | 0.0020509 | 1.4915 | 343.66 | 1859.83 | 0.04117 |
| TRINITY_DN3313835_c6_g2_i3 | 0.0027182 | 0.0029221 | 0.9302 | 51.29 | 1396.98 | 0.04425 |
| TRINITY_DN3417822_c3_g2_i2 | 0.0024484 | 0.0027053 | 0.9050 | 478.65 | 1458.13 | 0.04702 |

c. bloom vs. post-bloom

| | average | sd | ratio | ava | avb | cumulative sum |
|---|---|---|---|---|---|---|
| **TRINITY_DN3567435_c13_g1_i1** | **0.0085442** | **4.672e-03** | **1.829** | **7807.2** | **1381.03** | **0.01159** |
| TRINITY_DN3565854_c4_g3_i5 | 0.0056244 | 1.861e-03 | 3.022 | 11.80 | 4133.35 | 0.01922 |
| TRINITY_DN3531545_c4_g1_i13 | 0.0048821 | 1.976e-03 | 2.471 | 4963.8 | 1361.30 | 0.02585 |
| TRINITY_DN3383058_c4_g1_i10 | 0.0039800 | 2.688e-03 | 1.481 | 3964.9 | 1036.28 | 0.03125 |
| TRINITY_DN3561515_c5_g1_i4 | 0.0035883 | 2.845e-04 | 12.613 | 64.05 | 2689.11 | 0.03612 |
| TRINITY_DN3455354_c3_g3_i2 | 0.0031505 | 3.037e-04 | 10.376 | 24.96 | 2331.49 | 0.04039 |
| TRINITY_DN3531545_c4_g1_i11 | 0.0031379 | 1.345e-03 | 2.334 | 3757.6 | 1439.78 | 0.04465 |
| TRINITY_DN3565854_c4_g3_i2 | 0.0028920 | 7.801e-04 | 3.707 | 7.66 | 2126.66 | 0.04857 |
| TRINITY_DN3567090_c10_g1_i2 | 0.0028720 | 1.036e-03 | 2.772 | 2911.7 | 776.19 | 0.05247 |
| TRINITY_DN3417822_c3_g2_i2 | 0.0028412 | 1.986e-03 | 1.4303 | 2269.4 | 241.16 | 0.05633 |
